# Uncertainty in Deep Learning for EEG under Dataset Shifts

**DOI:** 10.1101/2025.07.09.663220

**Authors:** Mats Tveter, Thomas Tveitstøl, Christoffer Hatlestad-Hall, Hugo L. Hammer, Ira R. J. Hebold Haraldsen

## Abstract

**Objective:** As artificial intelligence (AI) is increasingly integrated into medical diagnostics, it is essential that predictive models provide not only accurate outputs but also reliable estimates of uncertainty. In clinical applications, where decisions have significant consequences, understanding the confidence behind each prediction is as critical as the prediction itself. Uncertainty modelling plays a key role in improving trust, guiding decision-making, and identifying unreliable outputs, particularly under dataset shift or in out-of-distribution settings. The primary aim of uncertainty metrics is to align model confidence closely with actual predictive performance, ensuring confidence estimates dynamically adjust to reflect increasing errors or decreasing reliability of predictions. This study investigates how different ensemble learning strategies affect both performance and uncertainty estimation in a clinically relevant task: classifying Normal, Mild Cognitive Impairment, and Dementia from electroencephalography (EEG) data.

**Approach:** We evaluated the performance and uncertainty of ensemble methods and Monte Carlo dropout on a large EEG dataset. The models were assessed in three settings: (1) in-distribution performance on a held-out test set, (2) generalisation to three out-of-distribution datasets, and (3) performance under gradual, EEG-specific dataset shifts simulating noise, drift, and frequency perturbation.

**Main results:** Ensembles consisting of multiple independently trained models, such as deep ensembles, consistently achieved higher performance in both the in-distribution test set and the out-of-distribution datasets. These models also produced more informative and responsive uncertainty estimates under various types of EEG dataset shifts.

**Significance:** These results highlight the benefits of ensemble diversity and independent training to build robust and uncertainty-aware EEG classification models. The findings are particularly relevant for clinical applications, where reliability under distribution shift and transparent uncertainty are essential for safe deployment.

## 1. Introduction

Recent advances in artificial intelligence (AI) have significantly enhanced its capabilities, establishing it as a critical component in various scientific and industrial applications. One of the fields where this technology has the greatest potential for impact is medical diagnostics and risk assessment, with predictive algorithms, research methodologies, and data analysis [1, 2]. Deep learning (DL), a subfield of AI [3], is notable for the depth and complexity of its models, which enables them to learn directly from raw data. These models often contain millions to billions of parameters and have achieved state-of-the-art performance across a wide range of tasks. In low-stakes or non-critical settings, such complexity is generally unproblematic, as occasional prediction errors are unlikely to have serious consequences. However, in clinical medicine, where predictions can have significant consequences for individuals, this complexity introduces critical challenges and safety concerns [4]. The potential for inaccurate predictions in high-stakes settings requires extensive evaluation and validation to ensure safe and effective integration of these models into clinical practice [5]. Two main strategies are commonly used to address the challenges of complexity and trust in AI; explainability and uncertainty modelling. Explainability methods aim to improve interpretability by clarifying the decision-making processes of AI models [6, 7], but do not assess the reliability of the decision [8] and are vulnerable to overconfident predictions and inputs specifically designed to mislead the model, known as adversarial attacks [9]. On the other hand, uncertainty modelling aims to inform users about the confidence of predictions, thereby enhancing the trustworthiness of the system [9].

For AI models to be trusted in clinical settings, they must not only be accurate but also reliably indicate when their predictions may be unreliable. Such uncertainty can stem from the inherent noise of biological signals and patient variability, or from the model’s own knowledge gaps due to limited training data [10, 11]. In practice, these challenges manifest as predictive uncertainty, which can be classified into three main types [5]: in-domain uncertainty for data drawn from the same distribution as the training set [12]; domain shift uncertainty, where the data is shifted from the training distribution [13, 14]; and out-of-distribution (OOD) uncertainty, where the data originate from an unknown distribution [15]. Effectively managing data shifts and OOD data is critical to ensure robust and safe deployment [16]. To achieve this, methods like Bayesian inference and ensembles are used to create well-calibrated models, where the output probability distribution aligns with the true likelihood of outcomes, making the model’s confidence explicitly interpretable [17]. The challenges of uncertainty are especially pronounced in neurophysiology, where signals are notoriously complex and noisy.

Electroencephalography (EEG) is a prime example, which is a non-invasive method for measuring the brain’s electrical activity from scalp electrodes. The recorded signals are multivariate time series that reflect the summation of postsynaptic potentials from neuronal populations, shaped by the orientation and proximity of the underlying signal sources [18]. However, it is highly susceptible to intrinsic and extrinsic artefacts, with a low signal-to-noise ratio [19]. DL and machine learning have been successfully applied to EEG data for tasks such as brain-computer interfaces, sleep staging, routine EEG interpretation [20] and prediction of cognitive states [19]. Uncertainty estimation in the context of EEG and deep learning has been proposed for several purposes. It has been used to assist in artefact removal from EEG signals [21] and to improve performance in brain–computer interfaces [22]. In seizure detection, uncertainty estimation has been applied to increase confidence in patient-level predictions [23]. In the domain of sleep staging, expressing uncertainty as a singular metric has been shown to enhance model transparency [24, 25], with a significant correlation observed between uncertainty estimates and model performance [24]. [26] extends this further by addressing uncertainty in motor-imagery tasks and aim to reduce uncertainty with the adaptive data augmentation approach [26]. Furthermore, previous work also demonstrated the use of uncertainty estimation in sex classification from EEG, highlighting how ensemble-based methods can improve predictive performance [27]. These applications demonstrate that quantifying uncertainty is a critical step toward developing more robust and trustworthy AI systems for clinical neurology.

In this study, we investigate how different ensemble strategies impact both predictive performance and uncertainty estimation in the classification of Normal, Mild Cognitive Impairment (MCI), and Dementia from resting-state EEG recordings. While grounded in a clinically relevant classification task, this work focuses on methodological evaluation of model performance and uncertainty. Our focus is on assessing model behaviour under realistic clinical conditions, including heterogeneous populations, real-world EEG variability, and distributional shifts. Specifically, we evaluate ensemble methods, including independently trained deep ensembles, checkpoint-based approaches, and methods such as Monte Carlo dropout and Stochastic Weight Averaging Gaussian, under three conditions: (1) a held-out in-distribution test set, (2) three relevant OOD datasets, and (3) controlled, EEG-specific dataset shifts constructed by systematically altering signal characteristics.

Through this comprehensive evaluation, we aim to answer two key questions: (1) which ensemble methods yield the most reliable predictions and well-calibrated uncertainty estimates across different datasets, and (2) how uncertainty relates to predictive performance across increasing levels of dataset shift. By examining these questions in the context of clinically relevant EEG classification, this study contributes to the development of more robust and trustworthy AI systems for clinical decision support.

## 2. Methods and materials

### 2.1 Dataset

#### 2.1.1 Training dataset

The EEG dataset used is the South Korean CAUEEG dataset [28], which includes 1,176 subjects clinically classified into three categories: normal, MCI and dementia. The dataset is pre-divided into 80% training, 10% validation, and 10% test sets, a split adopted in this study for consistency and comparability. The cohort had an age range of 23-96 years (mean = 71.13 ± 9.81). The dataset consists of EEG recordings in resting state, with segments associated with photic stimulation. Only sections preceding the photic stimuli were analysed, starting from the first eyes-closed marker. The preprocessing pipeline included bandpass filtering between 1.0 Hz and 45.0 Hz, re-referencing to the average, and removing line noise using a notch filter. EEG data were segmented into non-overlapping 30-second epochs. Autoreject [29] was then applied to remove epochs with excessive noise and artefacts and to repair channels. To improve model generalisation, data augmentation techniques were applied, limited to those that did not overlap with dataset shift, including time reversal, sign flipping, and smooth time masking (mask length = 20). Four epochs were selected per subject, ensuring maximal spread across the available epochs to increase variance. The final prediction for each subject was obtained by averaging the softmax outputs of the predictions across the four epochs.

In addition to EEG, the subject’s age was also used as input to the model due to its impact on performance [28], since age is a significant predictor of dementia [30]. Age data were normalised using min-max scaling, and random Gaussian noise (*µ* = 0, *σ* = 0.2) was added with a probability of 0.5 to account for variability and improve model generalisation.

#### 2.1.2. OOD datasets

Three OOD datasets were used: Militadous [31]‡, MPI [32], and TDBRAIN [33]. All OOD datasets were subjected to the same pre-processing pipeline as CAUEEG, and only the resting state EEG data were selected. First, a subset of channels was selected to match the 19 channels in the CAUEEG. In TDBRAIN and MPI, channels T7, T8, P7 and P8 were renamed to account for the absence of channels T3, T4, T5 and T6, as they cover the same regions, reflecting differences in naming conventions between the 10–20 and 10–10/10–5 system. Additionally, in MPI, some subjects were excluded due to missing electrodes. The labels varied between the datasets. The Miltiadous dataset comprises 88 participants (44 female, 44 male) aged 49–79 years (mean = 66.17 *±* 7.32), clinically stratified into Alzheimer’s disease (*n* = 36), frontotemporal dementia (*n* = 24), and healthy controls (*n* = 30). For the current analysis, the two dementia groups were merged into a single ‘dementia’ class. The MPI dataset includes 131 healthy subjects (90 male, 41 female) aged 22.5–77.5 years (mean = 38.26 *±* 20.02), after excluding participants with missing channels. The TDBRAIN dataset provided 27 healthy control participants (15 male, 12 female), aged 18.3–55.3 years (mean = 29.02 *±* 11.26), selected from a larger clinical cohort referred for neurophysiological evaluation. Only individuals with complete data were kept. In addition to EEG, age was also used as an input feature for the OOD datasets.

### 2.2 Designing dataset shifts

Multiple synthetic shifts were developed to simulate targeted perturbations in EEG data, enabling the evaluation of model performance and uncertainty under controlled distributional shifts. The objective was to introduce controlled, gradual shifts that preserve the complex signal integrity of EEG, in order to systematically evaluate the model’s response to specific types of signal alteration. This approach facilitates the evaluation of model robustness under varying degrees of dataset shift.

This framework was defined by a shift intensity parameter *x*. Notably, the interpretation of *x* depends on the type of dataset shift being applied. For static shifts, such as channel interpolation and channel rotation, the shift intensity *x* determined the proportion of EEG channels that were affected, with values defined as *x* ∈ {0.1, 0.25, 0.5, 0.75, 0.9, 1.0}. In contrast, for magnitude-based shifts, such as additive Gaussian noise, the parameter *x* specified the severity of the perturbation, for example by controlling the standard deviation of the noise applied.

To formalise the notation, let *S*_*i*_(*t*) represent the signal of channel *i* at time *t*. Importantly, for the magnitude-based shifts, the noise applied to each channel is sampled independently, ensuring that the variations are channel-specific rather than global. The noise magnitude for dataset shifts is drawn from the distribution:

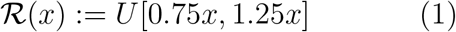

where *U* [*a, b*] is the uniform distribution in the interval [a,b] and *x* specifies the intended noise magnitude. This introduces moderate variability around *x*, producing more heterogeneous but bounded perturbations. In some shift types, the direction of the noise was also randomised, represented by:

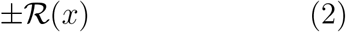

This represents a 50% probability of positive or negative perturbation per instance. Determining an appropriate value for *x* is non-trivial. For each type of shift, several values of *x* were empirically tested to explore their effect on the signal. Some of the implemented shift types have previously been proposed as data augmentation strategies [34].

#### 2.2.1. Gaussian noise

In EEG recordings, noise can originate from multiple sources, including sensor defects, environmental interference, or physiological artefacts (e.g. muscle activity). Let *ϵ*_*i*_(*t*) denote Gaussian noise sampled from a normal distribution 𝒩(0, *σ*^2^). The augmented signal 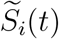 is defined by

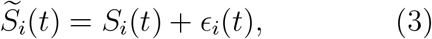

#### 2.2.2. Interpolation

To simulate sensor dropout or channel malfunction, a subset of EEG channels was temporarily removed and then reconstructed using interpolation based on a Minimum Norm Estimate [35]. At maximum shift intensity, all channels except two (Cz and Pz) are removed and interpolated.

#### 2.2.3. Bandstop filtering

The signals were bandstop filtered to remove specific frequency bands. The following frequency bands were investigated: delta (1–4 Hz), theta (4–8 Hz), alpha (8–12 Hz) and beta (12–30 Hz). Finally, a low-pass filter, filtering out gamma frequencies (>30Hz). This shift is the only shift which does not have a shift intensity.

#### 2.2.4. Channel shuffling

To simulate variations in electrode placement, a subset of EEG channels was randomly shuffled. The selected channels are randomly reordered, with no channel remaining in its original position, and the same permutation is applied to all subjects. This shift introduces controlled spatial variability while preserving the temporal structure of the signals.

#### 2.2.5.Amplitude modulation

To simulate the variability in signal gain and electrode sensitivity, the amplitude of each EEG channel was modulated by a scalar factor. Let *k*_*i*_ ~ ℛ(*k*) be a modulation factor of channel *i*.

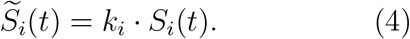

#### 2.2.6. Phase shift

To introduce variability into the dynamics of EEG signals, we applied a phase shift to each channel. Let *ϕ*_*i*_ ~ *±* ℛ(*ϕ*) be the phase change in radians, and let ℋ{*S*_*i*_(*t*)} denote the Hilbert transform of *S*_*i*_(*t*).

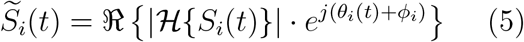

#### 2.2.7. Circular shift

To introduce variability in the temporal alignment of the EEG signals, we applied a circular change to each channel, where *k*_*i*_ ~ ± ℛ(*k*), and *T* is the total number of time samples.

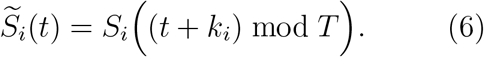

#### 2.2.8. Slow drift

In order to simulate slow-varying offset fluctuations (e.g., due to perspiration or electrode impedance changes), a slow drift was added to each EEG channel. The drift function Δ_*i*_(*t*) is constructed as:

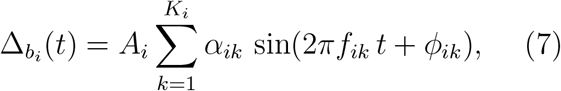

where *A*_*i*_ ~ ℛ(𝒜) is the maximum amplitude and a *K*_*i*_ ~ ℛ (𝒦) is the number of sinusoidal waves, *α*_*ik*_ ~ *U* [0.5, 1.0] is a weighting factor for the *k*-th sinusoid, *f*_*ik*_ ~ *U* [0.01, 0.2] is the frequency and *ϕ*_*ik*_ ~ *U* [0, 2*π*] is the phase of each sinusoid. The drifted signal is then defined as:

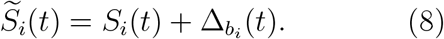

#### 2.2.9. Non-linear time warping

To introduce temporal distortions in EEG signals, a nonlinear time-warping transformation was applied. We define the warp function *w*(*t*) as:

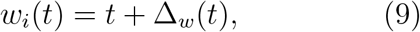

where 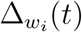 is a sinusoidal displacement:

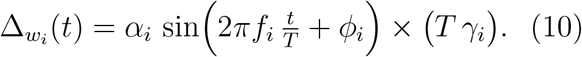

Here, *α*_*i*_ ~ ℛ(*α*), *T* is the total number of time points *f*_*i*_ ~ *U* [0.01, 0.2] is the random frequency, *ϕ*_*i*_ ~ *U* [0, 2*π*] is a random phase change and *λ*_*i*_ ~ [0.5, 1.0] is a randomly sampled scaling factor. Since *w*_*i*_(*t*) often yields non-integer indices, the final warped signal 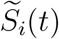 is obtained by linearly interpolating *S*_*i*_(*t*) at these warped indices:

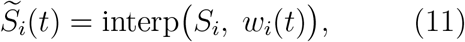

#### 2.2.10. Peak shift

To simulate a shift in the alpha band (8-12 Hz), we first isolate the alpha component *α*_*i*_(*t*) from an EEG signal *S*_*i*_(*t*) using band-pass filtering. We define the residual outside the alpha band as:

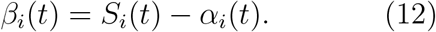

Next, we compute the analytic signal of *α*_*i*_(*t*) using the Hilbert transform:

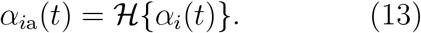

Then a frequency shift 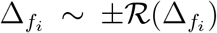 was applied to the analytic signal by multiplying it with a complex exponential and taking the real part:

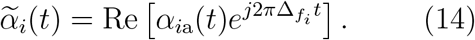

Finally, we reconstruct the shifted EEG signal by adding 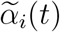 back to the residual *β* (*t*):

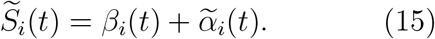

### 2.3 Model

The primary DL model used in this study is the InceptionNetwork [36], which has shown strong performance in EEG classification tasks [37, 38, 27, 39]. All ensembles were constructed using this single architecture, and for ensemble types that require repeated training runs, the ensemble size was fixed at five, consistent with previous studies [40]. For ensemble methods requiring only a single training run, the ensemble size was set to 20 to balance predictive performance and computational cost. Hyperparameter tuning was performed using the Optuna framework [41], employing the Tree-structured Parzen Estimator (TPE) algorithm [42]. A total of 125 trials were conducted per ensemble type to ensure fairness between the ensemble methods. The randomly initialised weight ensemble was excluded from tuning due to its lack of adjustable parameters.

#### Base model

The base model consists of a six-layer CNN with 20 units per layer and a maximum kernel size of 60. It includes two fully connected layers with 256 and 128 neurons, employing batch normalisation and dropout (rate: 0.5) to enhance generalisation. The ensemble specific parameters and changes in the model can be found in the github repository *§*.

#### Temperature scaling

This study employed temperature scaling, a post-hoc calibration technique proposed by Guo et al. [43], which rescales the logits by a scalar temperature parameter to adjust the confidence of the softmax outputs without altering the predicted class labels. Temperature scaling was applied to the aggregated logits of the ensemble, and the temperature values were optimised using the validation set.

#### 2.3.1. Deep Ensemble

The Deep Ensemble [40] is trained by initialising multiple models independently using different random seeds, allowing each model to explore distinct regions of the parameter space.

#### 2.3.2. Bagging Ensemble

The Bagging Ensemble [44] approach works by randomly drawing a subset of the training set with replacement and using this subset to train a classifier. By repeating this process multiple times, an ensemble of classifiers trained on different subsamples of the data is created. This ensemble used smaller versions of the base models as it is trained using less data.

#### 2.3.3. Depth Ensemble

The Depth Ensemble takes advantage of architectural variation by altering the number of inception modules within the InceptionNetwork. By varying the number of Inception modules, the Depth Ensemble creates models with different capacities to extract and represent features.

#### 2.3.4. Augmentation Ensemble

The augmentation ensemble technique differs from the deep ensemble by also using different data augmentation techniques per model in the ensemble.

#### 2.3.5. Monte Carlo Dropout

The Monte Carlo dropout (MCD) [45] approximates a Bayesian method by randomly dropping neurons during inference time. This *sampling* with multiple forward passes of different architectures with different neurons turned off generates a computationally efficient ensemble. In this study, 50 stochastic forward passes were used during inference.

#### 2.3.6 Snapshot Ensemble

The snapshot ensemble [46] generates *M* models in a single training session. This method rapidly changes the learning rate using a cyclic cosine annealing schedule [47]. This approach enables the model to explore multiple local minima, with the schedule periodically increasing the learning rate to escape the current minima.

#### 2.3.7. Stochastic Weight Averaging Gaussian

Stochastic Weight Averaging Gaussian (SWAG) [48] extends Stochastic Weight Averaging (SWA) [49] by modeling the distribution of weights in parameter space. Instead of averaging weights at selected checkpoints, SWAG estimates a Gaussian distribution over the weights by collecting the mean and lowrank covariance from multiple points during training. In our implementation, weights were collected from a partially trained model to estimate the SWAG distribution. At test time, 50 weight samples were drawn to form an ensemble, and their predictions were averaged.

#### 2.3.8. Fast Geometric Ensemble

The Fast Geometric Ensemble (FGE) [50] technique is similar to the snapshot ensemble [46] in the way models are saved during one iteration of the training and similar to SWA by starting with a pre-trained model. It differs from SE by using a piecewise linear cyclical learning rate schedule rather than the cosine schedule, and, in addition, the cycles are usually smaller in magnitude.

#### 2.3.9. Ensemble taxonomy

The ensemble methods can be grouped into three categories based on how they create and maintain diversity. Multi-run ensembles, including Deep Ensemble, Bagging, Augmentation, and Depth, train multiple independently initialised models. Checkpoint-based methods, such as Snapshot and FGE, extract ensemble members from intermediate stages of a single training run. Sampling-based methods, including MCDropout and SWAG, derive diversity by sampling during inference using a single trained model in the case of MCD and drawing weights from a learnt posterior approximation in the case of SWAG.

### 2.4. Performance metrics

For this study, Area-Under-the-curve (AUC) [51] is used to compare the performance of the models. The AUC has a value between 0.0 and 1.0, with 0.5 representing random guessing. For multiple classes, the One-vs-Rest (Ovr) strategy is used, where the AUC is computed for each class against all others and then averaged using a weighted approach to provide an overall performance measure. For per-class AUC, Ovr is applied without weighting, where AUC is calculated separately for each class. For comparisons involving OOD datasets with fewer classes, recall and precision are reported per class. Recall reflects the proportion of correctly identified positives, while precision indicates the accuracy of positive predictions.

### 2.5. Uncertainty metrics

*Brier score* The Brier score [52] is a strictly proper scoring rule that evaluates the quality of probabilistic predictions by jointly capturing both accuracy and calibration. It measures how close the predicted probability distribution is to the actual outcome. A lower score reflects better performance, indicating that the model not only predicts the correct class more often but also assigns appropriate levels of confidence to its predictions. In other words, the Brier score penalizes both incorrect predictions and over- or underconfident probability estimates. A perfectly calibrated and accurate model would achieve a score of 0. The Brier score for multiclass classification is defined as:

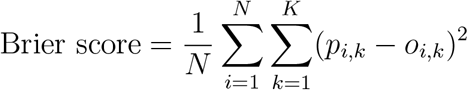

where *N* is the number of instances, *K* is the number of classes, *p*_*i,k*_ is the predicted probability for instance *i* belonging to class *k* and *o*_*i,k*_ is the actual outcome which is 1 if the instance *i* belongs to class *k*, and 0 otherwise.

*Expected Calibration Error (ECE)* The ECE quantifies how well the predicted probabilities match the observed frequencies, providing information on model calibration. Predictions are grouped into bins *B* according to their confidence scores. For each bin *b*, the accuracy and confidence are calculated as:

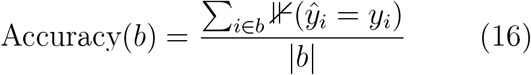

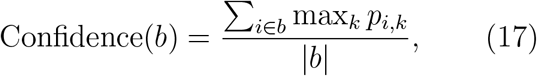

where max_*k*_ *p*_*i,k*_ is the highest predicted probability for sample *i*, and ⊮ (*ŷ*_*i*_ = *y*_*i*_) is an indicator function that equals 1 if the predicted class *ŷ*_*i*_ = arg max_*k*_ *p*_*i,k*_ matches the true label*y*_*i*_, and 0 otherwise. The ECE is then calculated as the weighted average of the absolute difference between accuracy and confidence across bins:

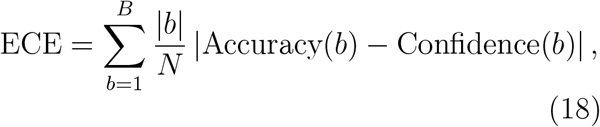

where |*b*| is the number of predictions in bin *b*, and *N* is the total number of samples. A lower ECE indicates better calibration, meaning that the predictions align more closely to the observed outcomes. The number of bins was set to 10.

## 3. Results

### 3.1. In-distribution test performance

Table 1 reports several metrics evaluating the performance of the different ensemble methods on the test set. Starting with overall AUC, we observe that the multi-run ensembles, including MCD, demonstrate strong predictive performance. In contrast, checkpoint ensembles perform significantly worse, with nonoverlapping confidence intervals. SWAG falls in between these groups. For accuracy, a metric more sensitive to class imbalance, all multirun models achieve over 60%, while checkpoint and sampling models fall below this threshold. Notably, SWAG displays high run-to-run variability. Looking at the per-class AUC, all models perform best in classifying normal and dementia cases, with the top AUC values exceeding 0.90. However, MCI classification remains challenging, with the best AUC reaching only 0.71.

**Table 1:** Performance on the holdout test set with 117 subjects, in predicting between the classes: Normal, MCI and Dementia. The performance is calculated across 5 runs per ensemble with mean and 95% confidence interval. Where N=Normal, M=MCI and D=Dementia for the 3 right columns.

| Ensemble Type | AUC | Accuracy (%) | Recall<sub>class</sub> | Precision<sub>class</sub> | AUC<sub>class</sub> |
| --- | --- | --- | --- | --- | --- |
| Deep Ensemble | $0.84 \pm 0.01$ | $67.18 \pm 2.07$ | N: $0.90 \pm 0.06$ | N: $0.71 \pm 0.07$ | N: $0.91 \pm 0.01$ |
| | | | M: $0.41 \pm 0.10$ | M: $0.59 \pm 0.02$ | M: $0.70 \pm 0.02$ |
| | | | D: $0.68 \pm 0.07$ | D: $0.69 \pm 0.06$ | D: $0.91 \pm 0.01$ |
| Bagging | $0.83 \pm 0.02$ | $65.13 \pm 3.93$ | N: $0.92 \pm 0.03$ | N: $0.68 \pm 0.06$ | N: $0.90 \pm 0.01$ |
| | | | M: $0.27 \pm 0.12$ | M: $0.56 \pm 0.12$ | M: $0.68 \pm 0.04$ |
| | | | D: $0.77 \pm 0.12$ | D: $0.66 \pm 0.05$ | D: $0.91 \pm 0.01$ |
| Depth | $0.83 \pm 0.02$ | $68.21 \pm 2.75$ | N: $0.92 \pm 0.06$ | N: $0.70 \pm 0.04$ | N: $0.91 \pm 0.02$ |
| | | | M: $0.40 \pm 0.12$ | M: $0.63 \pm 0.06$ | M: $0.69 \pm 0.03$ |
| | | | D: $0.71 \pm 0.08$ | D: $0.69 \pm 0.07$ | D: $0.90 \pm 0.01$ |
| Augmentation | $0.84 \pm 0.02$ | $67.69 \pm 2.30$ | N: $0.92 \pm 0.04$ | N: $0.68 \pm 0.04$ | N: $0.91 \pm 0.01$ |
| | | | M: $0.33 \pm 0.08$ | M: $0.62 \pm 0.07$ | M: $0.71 \pm 0.04$ |
| | | | D: $0.78 \pm 0.06$ | D: $0.71 \pm 0.04$ | D: $0.92 \pm 0.02$ |
| MCD | $0.80 \pm 0.03$ | $59.66 \pm 5.69$ | N: $0.86 \pm 0.12$ | N: $0.65 \pm 0.10$ | N: $0.87 \pm 0.02$ |
| | | | M: $0.29 \pm 0.15$ | M: $0.49 \pm 0.08$ | M: $0.66 \pm 0.02$ |
| | | | D: $0.61 \pm 0.37$ | D: $0.66 \pm 0.13$ | D: $0.89 \pm 0.04$ |
| Snapshot | $0.71 \pm 0.03$ | $52.82 \pm 5.59$ | N: $0.64 \pm 0.11$ | N: $0.65 \pm 0.02$ | N: $0.80 \pm 0.02$ |
| | | | M: $0.39 \pm 0.06$ | M: $0.40 \pm 0.08$ | M: $0.53 \pm 0.04$ |
| | | | D: $0.56 \pm 0.07$ | D: $0.53 \pm 0.05$ | D: $0.80 \pm 0.02$ |
| SWAG | $0.77 \pm 0.04$ | $40.00 \pm 16.46$ | N: $0.38 \pm 0.52$ | N: $0.76 \pm 0.22$ | N: $0.86 \pm 0.04$ |
| | | | M: $0.16 \pm 0.43$ | M: $0.28 \pm 0.54$ | M: $0.58 \pm 0.05$ |
| | | | D: $0.74 \pm 0.54$ | D: $0.30 \pm 0.30$ | D: $0.87 \pm 0.03$ |
| FGE | $0.74 \pm 0.01$ | $57.91 \pm 4.48$ | N: $0.76 \pm 0.03$ | N: $0.67 \pm 0.04$ | N: $0.84 \pm 0.01$ |
| | | | M: $0.37 \pm 0.07$ | M: $0.43 \pm 0.07$ | M: $0.56 \pm 0.02$ |
| | | | D: $0.58 \pm 0.07$ | D: $0.58 \pm 0.04$ | D: $0.83 \pm 0.01$ |

For the Normal class, the recall of multirun ensembles is high (≥ 0.90), moderately high for Dementia (≥ 0.68), and low for MCI (0.27–0.41). Precision follows the exact ordering, with about 0.70 for Normal and Dementia and around 0.60 for MCI. Checkpoint ensembles show markedly lower recall for Normal, reduced recall for Dementia, and similar recall for MCI; their precision is comparable to the multi-run group for Normal but lower for both Dementia and MCI. MCD confidence intervals overlap those of the multi-run models in every class, yet its average recall for Normal and Dementia is slightly lower, and the intervals themselves are wider. For precision, MCD has a similar performance to that of multi-run ensembles. SWAG displays the greatest instability, its recall, and precision intervals are extremely broad, with dementia recall dipping to near zero in some runs and reaching 1.0 in others, underscoring substantial run-to-run variability.

### 3.2. Out-of-Distribution

#### 3.2.1. Performance

Table 2 presents class-wise recall and precision for three OOD cohorts. In Militadous, the multi-run ensembles maintain strong recall for the normal class (0.82-0.89) and moderate recall for dementia (0.52-0.57). Their precision remains balanced, about 0.50 for Normal and above 0.93 for Dementia. Depth is slightly below the average of the other multi-run models, but its confidence intervals overlap. MCD shows similar behaviour for Normal but lower and more variable recall for Dementia, with precision similar to the multi-run ensembles. The checkpoint ensembles lose much more recall and precision, and SWAG shows the widest run-to-run variability, making its performance unreliable.

**Table 2:**
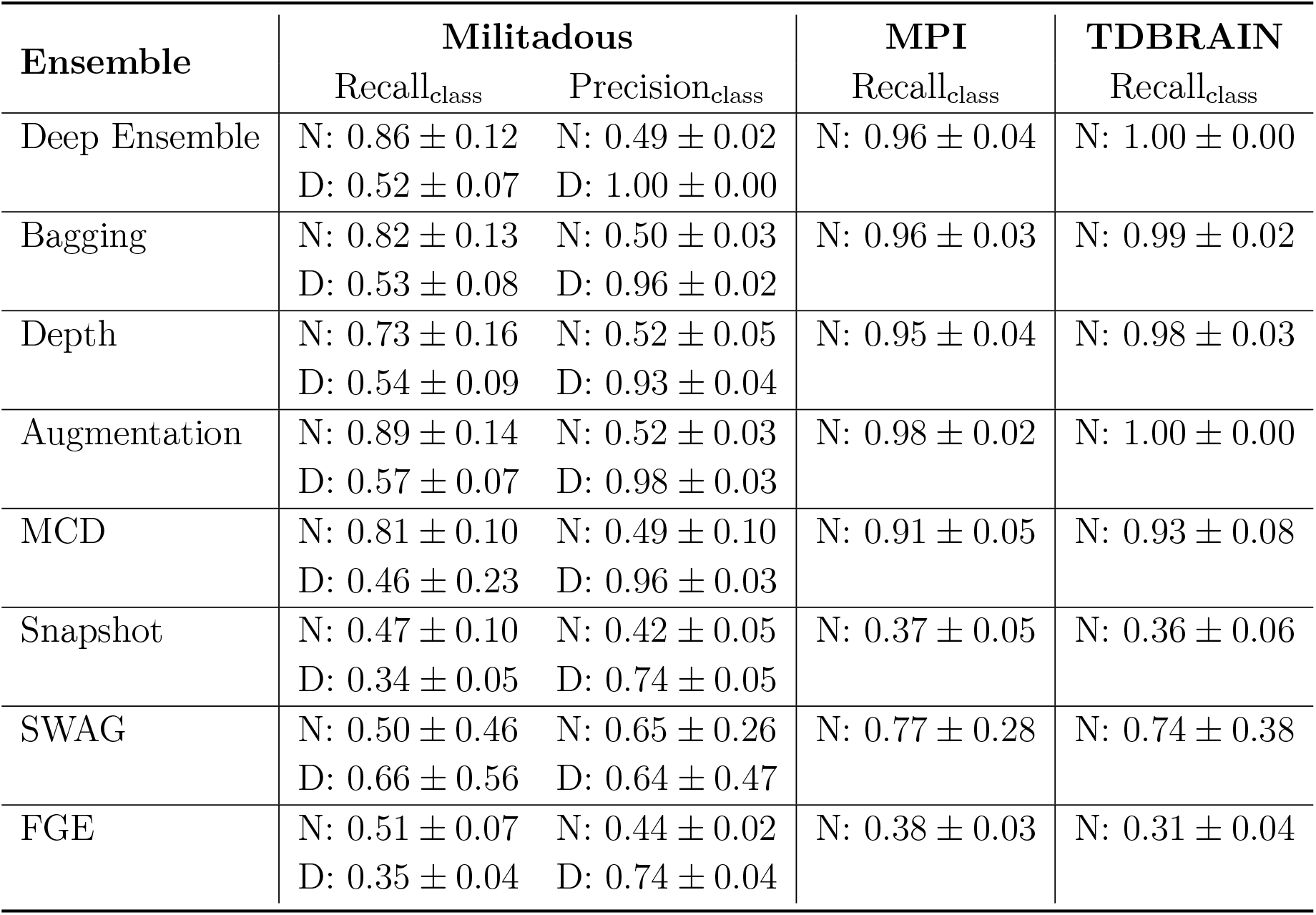
Comparison of precision and recall calculated per class in the 3 OOD datasets. The performance is calculated across 5 runs per ensemble with mean and 95% confidence interval. Where N=Normal and D=Dementia.

The single-class cohorts MPI and TD-BRAIN follow the same ranking pattern. All multi-run methods recall almost every normal subject (about 0.95-1.00). MCD is slightly lower on average, but its intervals overlap. Checkpoint ensembles miss a substantial share of normals, and while SWAG average recall is higher, its confidence intervals are so wide that its reliability remains doubtful.

#### 3.2.2. Uncertainty calibration

Table 3 summarises the Brier score and ECE for each ensemble on the test set and three OOD cohorts. On the test set, the multi-run ensembles record the lowest Brier values (0.45–0.47) and mid-range ECE error (0.10–0.13). The MCD Brier score is slightly higher and its ECE is slightly lower, but both confidence intervals overlap those of the multi-run group. The checkpoint ensembles have noticeably higher Brier scores than the multi-run methods, while their ECE is on average lower, though again with overlapping intervals. SWAG stands out with the highest Brier and ECE values and the widest confidence intervals. In the Militadous cohort, every ensemble shows a higher Brier score than on the test set, except SWAG, but with much wider confidence intervals, and the Augmentation ensemble, which remains stable (0.45 to 0.46). ECE error is largely stable for the multi-run methods with ECE near 0.10. In contrast, the checkpoint and sampling methods all see noticeably higher ECE values and broader intervals than on the test set. In the single-class MPI and TDBRAIN cohorts, Brier scores decrease for the multi-run ensembles and the sampling methods relative to the test set, whereas the checkpoint ensembles exhibit higher Brier scores. ECE values increase for every ensemble in both cohorts relative to their test set values.

**Table 3:** Uncertainty comparison of ensemble methods (mean ±95% CI over 5 runs) calculated across the test set and three OOD datasets.

| (a) Test (Normal/MCI/Dementia) |  |  | (b) Militadous (Normal/Dementia) |  |  |
| --- | --- | --- | --- | --- | --- |
| Ensemble | Brier | ECE | Ensemble | Brier | ECE |
| Deep Ensemble | $0.45 \pm 0.01$ | $0.10 \pm 0.02$ | Deep Ensemble | $0.52 \pm 0.02$ | $0.10 \pm 0.03$ |
| Bagging | $0.47 \pm 0.01$ | $0.11 \pm 0.03$ | Bagging | $0.50 \pm 0.06$ | $0.08 \pm 0.02$ |
| Depth | $0.47 \pm 0.00$ | $0.13 \pm 0.06$ | Depth | $0.54 \pm 0.05$ | $0.11 \pm 0.05$ |
| Augmentation | $0.45 \pm 0.01$ | $0.10 \pm 0.04$ | Augmentation | $0.46 \pm 0.03$ | $0.10 \pm 0.03$ |
| MCD | $0.51 \pm 0.05$ | $0.08 \pm 0.02$ | MCD | $0.58 \pm 0.15$ | $0.14 \pm 0.07$ |
| Snapshot | $0.57 \pm 0.03$ | $0.08 \pm 0.05$ | Snapshot | $0.75 \pm 0.04$ | $0.18 \pm 0.08$ |
| SWAG | $0.64 \pm 0.08$ | $0.17 \pm 0.08$ | SWAG | $0.53 \pm 0.13$ | $0.24 \pm 0.07$ |
| FGE | $0.55 \pm 0.01$ | $0.08 \pm 0.04$ | FGE | $0.75 \pm 0.01$ | $0.13 \pm 0.03$ |

Table 3: Uncertainty comparison of ensemble methods (mean $\pm 95\%$ CI over 5 runs) calculated across the test set and three OOD datasets.
| (c) MPI (Normal) |  |  | (d) TDBRAIN (Normal) |  |  |
| --- | --- | --- | --- | --- | --- |
| Ensemble | Brier | ECE | Ensemble | Brier | ECE |
| Deep Ensemble | $0.11 \pm 0.05$ | $0.17 \pm 0.02$ | Deep Ensemble | $0.06 \pm 0.03$ | $0.16 \pm 0.04$ |
| Bagging | $0.13 \pm 0.06$ | $0.20 \pm 0.04$ | Bagging | $0.08 \pm 0.02$ | $0.19 \pm 0.03$ |
| Depth | $0.18 \pm 0.08$ | $0.24 \pm 0.05$ | Depth | $0.14 \pm 0.06$ | $0.25 \pm 0.06$ |
| Augmentation | $0.11 \pm 0.05$ | $0.18 \pm 0.05$ | Augmentation | $0.06 \pm 0.03$ | $0.16 \pm 0.06$ |
| MCD | $0.24 \pm 0.09$ | $0.25 \pm 0.03$ | MCD | $0.26 \pm 0.10$ | $0.32 \pm 0.02$ |
| Snapshot | $0.71 \pm 0.02$ | $0.18 \pm 0.05$ | Snapshot | $0.70 \pm 0.02$ | $0.24 \pm 0.02$ |
| SWAG | $0.40 \pm 0.24$ | $0.24 \pm 0.15$ | SWAG | $0.42 \pm 0.25$ | $0.28 \pm 0.15$ |
| FGE | $0.73 \pm 0.01$ | $0.16 \pm 0.02$ | FGE | $0.72 \pm 0.02$ | $0.28 \pm 0.06$ |

### 3.3. Dataset shifts

#### 3.3.1. Performance and uncertainty calibration

We evaluated the models performance using AUC, and for uncertainty analysis, we investigated the Brier score and ECE.

Amplitude shifts (sub-figures 1a, 1b, and 1c) reveal three distinct patterns. Checkpoint ensembles maintain almost flat curves: their AUC, Brier score, and ECE remain stable across the full range of shift intensities. The sampling methods show worse performance, and MCD shows higher Brier and ECE at the lowest shift intensities. Multirun ensembles show the highest performance around (×0.001–×2), but their performance drops below the checkpoint models once the signal is amplified beyond about ×2. Between 0.25× and 2× scaling, the multi-run ensembles show lower Brier scores than the checkpoint models, but outside this window, their Brier scores are generally higher. Across the full amplitude range, they also exhibit higher ECE values and broader confidence intervals. Across all methods, the Augmentation ensemble achieves marginally better performance, although the differences are not statistically significant.

**Figure 1:**
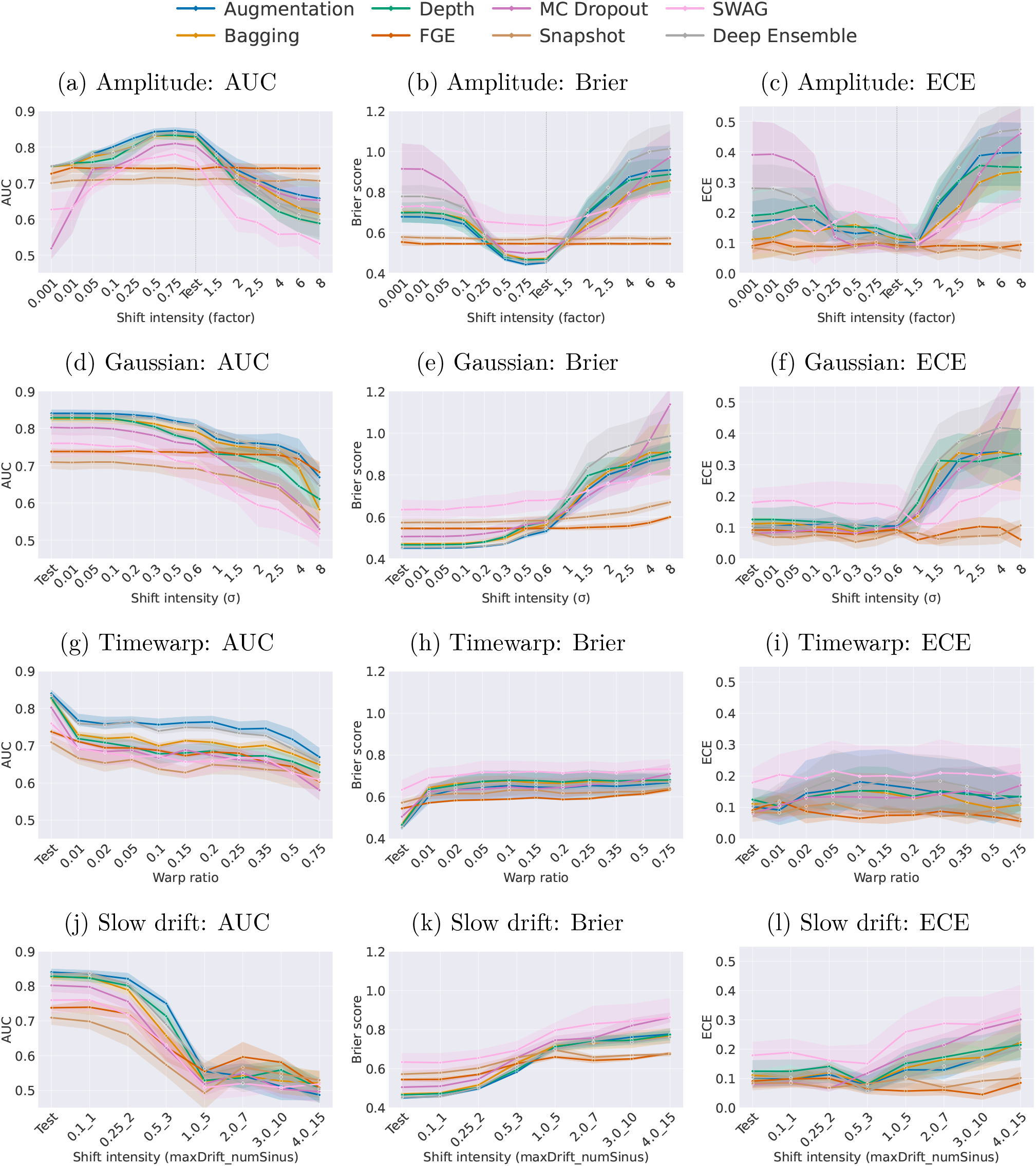
Illustrating ensemble performance, uncertainty and calibration under four dataset shifts. Columns plot three metrics, AUC, Brier score and ECE, against shift intensity. The rows (top to bottom) represent shift types: (1) amplitude scaling (signal multiplied by intensity), (2) Gaussian noise (added noise), (3) time warp (local compression/expansion), and (4) slow drift (added low-frequency sinusoidal waves). Ensemble methods are labelled in the top of the figure.

Under Gaussian noise (sub-figures 1d, 1e, and 1f) all ensembles maintain baseline AUC until the noise standard deviation reaches about *σ* = 0.2. Beyond this point, the multi-run methods continue to rank highest, with Augmentation and Deep ensembles remaining the best performers all the way to *σ* = 8 despite a gradual decline. They also maintain the lowest Brier scores up to roughly *σ* = 0.6. Checkpoint models are stable with FGE keeping both AUC and Brier score essentially flat until around *σ* = 4, whereas Snapshot performance declines at a rate similar to the multi-run ensembles, but with a stable Brier score. The sampling methods are initially lower in performance than multi-run but higher than checkpoint ensembles, following the multi-run curves in both AUC and Brier. The ECE is stable for all ensembles until about *σ* = 0.6 beyond that, ECE rises sharply for the multi-run and sampling methods while remaining nearly constant for the checkpoint models.

Timewarp shifts (sub-figures 1g, 1h, and 1i) result in an immediate and substantial drop in performance for all ensembles, even at the lowest shift intensity. Performance continues to decline as the shift increases for most ensembles, except Augmentation and Deep ensembles, which keep the performance at around 0.2. The Brier scores reflect this trend, with the largest increase occurring after the first shift, followed by a more gradual rise. The checkpoint ensembles, while starting with a slightly higher baseline Brier score, show a flatter trajectory across shift levels. ECE values remain relatively stable on average for all models, but with wider confidence intervals.

Slow drift (sub-figures 1j, 1k, and 1l) causes a gradual decline in performance below 0.25 Hz / 2 sinusoidal components, followed by a sharp drop between 0.25 Hz and 1.0 Hz / 5 sinus waves. While a larger grid of combinations was tested, only a representative subset is shown to highlight the overall trend. The augmentation ensemble seems to have a slower descent, at least until the mark of 1.0 Hz / 5 sinus waves. Beyond this point, model performance remains generally poor across all ensembles, with an AUC of around 0.5. The Brier scores follow a similar pattern, although the checkpoint models display a flatter curve after the 1.0 Hz / 5-wave mark. ECE values remain stable for most models up to 0.5 Hz / 3 sinus waves, after which calibration quality begins to degrade, again with checkpoint models maintaining lower ECE compared to the rest.

Phase shifts (sub-figures 2a, 2b and 2c) result in decreased performance across all shift levels, with the most pronounced drop occurring at phase offsets of *π/*2 and 3*π/*2 (i.e., 90° and 270°). The relative ranking of ensemble performance seen on the test set remains largely consistent across all phase shifts, with the augmentation and deep ensemble performing slightly better throughout the shift intensities. The Brier score follows a similar pattern, reflecting increased uncertainty at *π/*2 and 3*π/*2. In contrast, ECE remains relatively stable across most shifts and for all ensembles, with notable increases only at *π/*2 and 3*π/*2.

**Figure 2:**
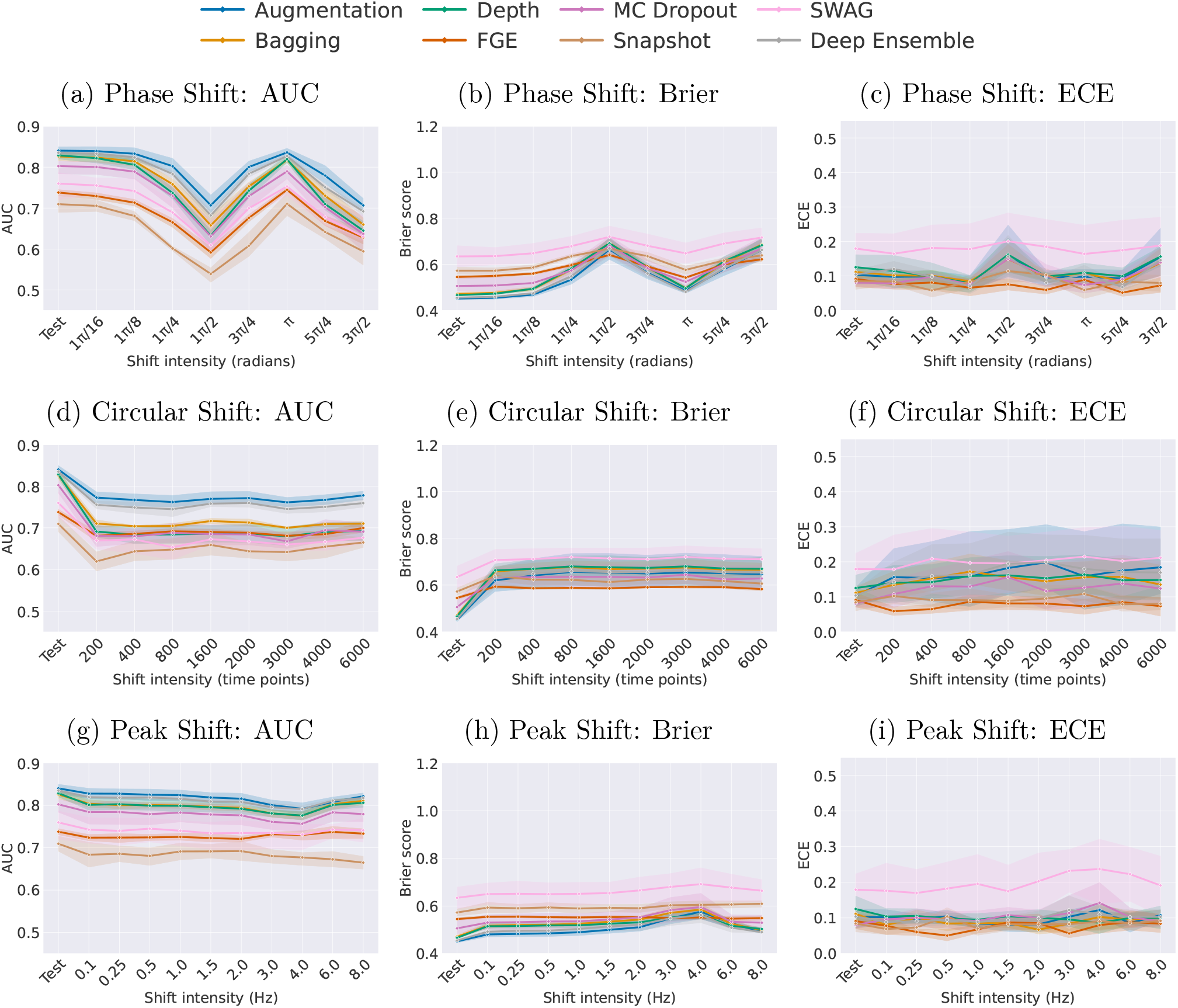
Illustrating ensemble performance, uncertainty and calibration under four dataset shifts. Columns plot three metrics, AUC, Brier score and ECE, against shift intensity. The rows (top to bottom) represent shift types: (1) phase shift (altering the signal’s phase), (2) circular shift (rotating the signal in time), and (3) peak shift (displacing the alpha-band peak). Ensemble methods are labelled across the top of each column.

Circular shifts (sub-figures 2d, 2e, and 2f) result in an immediate decline in performance across ensembles, with minor sensitivity to the magnitude of the shift. However, except for the baseline, the deep ensemble and augmentation ensemble perform significantly better than the rest. The Brier score shows an increase in uncertainty from the first shift level, followed by a relatively stable plateau. For ECE, the average calibration remains largely stable across all ensembles, although a slow increase is observed for some. Notably, the confidence intervals for ECE are wide.

Peak shifts (sub-figures 2g, 2h, and 2i) result in a gradual, slow decline in performance, with more noticeable drop occurring around the 2–4 Hz range and the relative ordering of the ensemble performance is kept throughout the shift. A similar pattern is observed in the Brier score, showing a slight increase between 2 and 6 Hz, except for the checkpoint ensembles. ECE remains relatively stable, on average, throughout the entire frequency range, with a slight increase at 4.0 Hz for some of the ensembles.

Interpolation (sub-figures 3a, 3b and 3c) show that the performance remains stable until approximately 75% of channels are removed, after which it begins to decline moderately, with a sharp drop beyond 90%. The relative ordering of the performance of the ensemble remains stable. The Brier score remains stable until approximately 90% of the channels are removed, at which point it increases sharply. A similar trend is observed for ECE.

**Figure 3:**
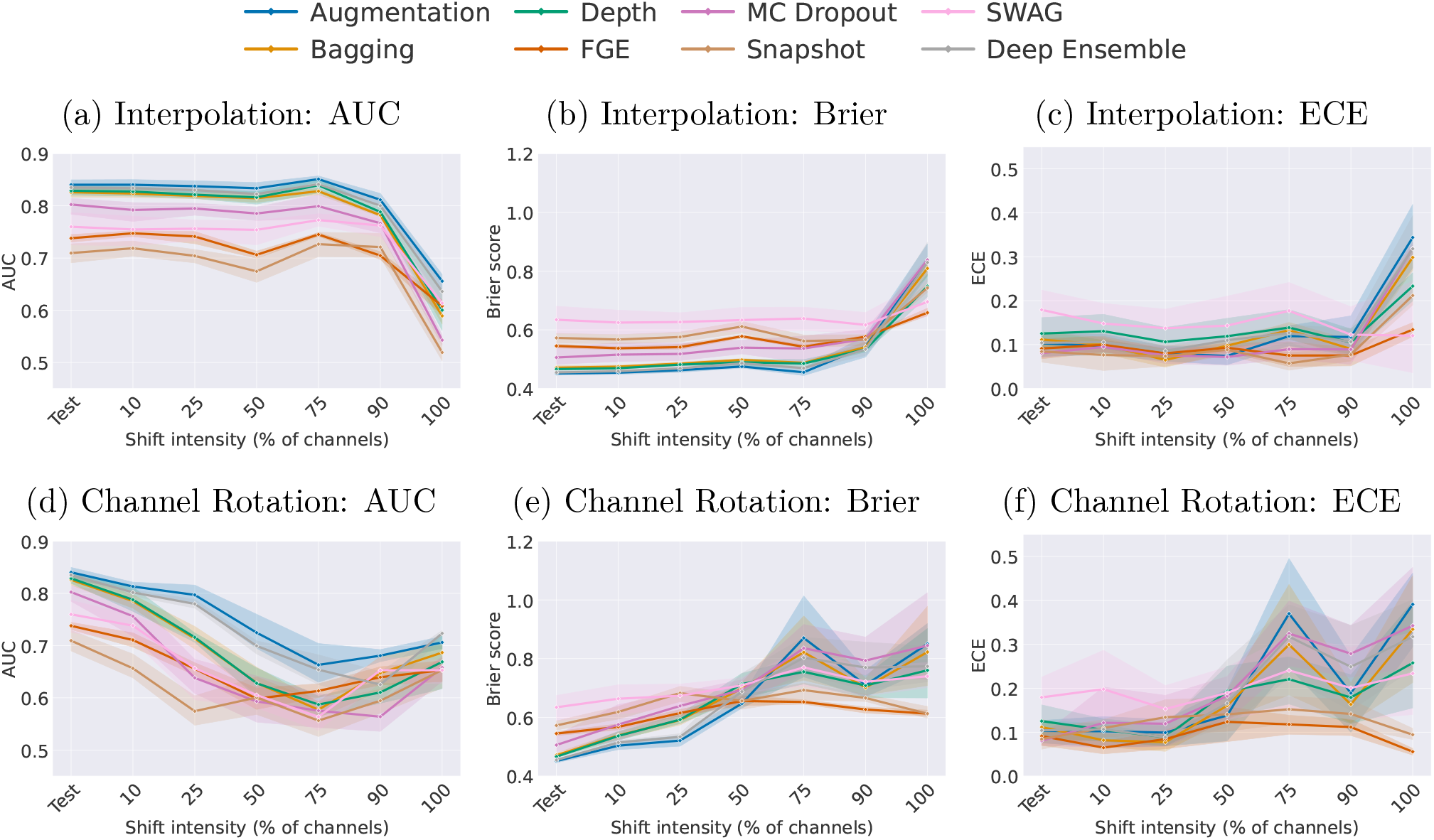
Illustrating ensemble performance, uncertainty and calibration under four dataset shifts. Columns plot three metrics, AUC, Brier score and ECE, against shift intensity. Rows (top to bottom) represent shift types: (1) interpolation (removing channels and interpolating), and (2) rotation (randomly rotating channel positions). Ensemble methods are labelled in the top of the figure.

Channel rotation (sub-figures 3d, 3e, and 3f) leads to a steady decline in performance across all ensembles. The Deep and Augmentation ensembles start higher and decline more slowly than the others until rotation reaches roughly 75–90 percent. Their Brier curves rise at a correspondingly slower rate. Most ensembles exhibit a sharp increase in Brier score and ECE once rotation exceeds about 75 percent, whereas the checkpoint models retain slightly better calibration in that range. ECE remains nearly unchanged for all methods up to 50 percent rotation, and beyond this point the checkpoint ensembles continue to hold the most stable ECE values.

Bandstop shifts (sub-figures 4a, 4b, and 4c) show that SWAG maintains stable, with larger confidence intervals, across all frequency bands. In contrast, the other ensembles exhibit a significant decline in the theta band, and all except MC Dropout and Snapshot also show a drop in performance when the delta band is removed. Additionally, Bagging, Depth, MC Dropout, Snapshot, and Deep Ensemble show reduced performance in the beta band. These patterns are mirrored in the Brier scores, with clear increases in the delta, theta, and beta ranges. ECE remains relatively stable across all frequency bands and ensemble types.

**Figure 4:**
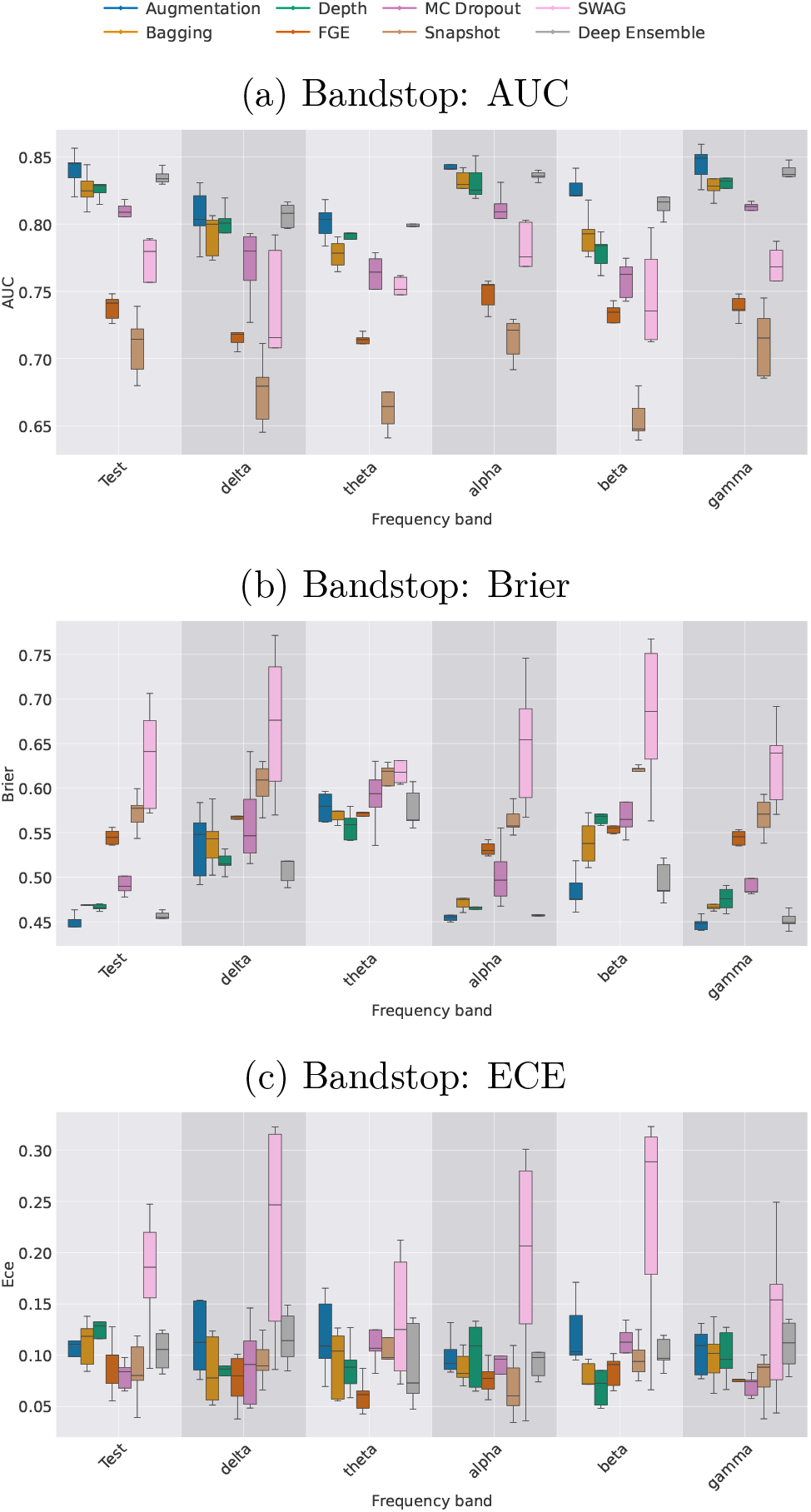
Performance, uncertainty and calibration of ensemble type to bandstop filtering. Rows plot, from top to bottom, (a) AUC, (b) Brier score and (c) ECE as a function of the specific frequency band removed. Ensemble methods are labelled in the top of the figure.

### 3.4. Per-Class

Figure 5 shows the AUC for the normal class (Figure 5a) and the dementia class (Figure 5b) when specific frequency bands are removed. For the Normal class, removing either the theta or beta bands results in a marked drop in AUC, indicating that these bands contribute to the normal class classification. The dementia class displays a partially overlapping but distinct pattern: theta-band removal again leads to the largest performance drop across methods and larger than for the normal class, while delta and beta removal have a more modest impact, particularly in the augmentationbased ensemble.

**Figure 5:**
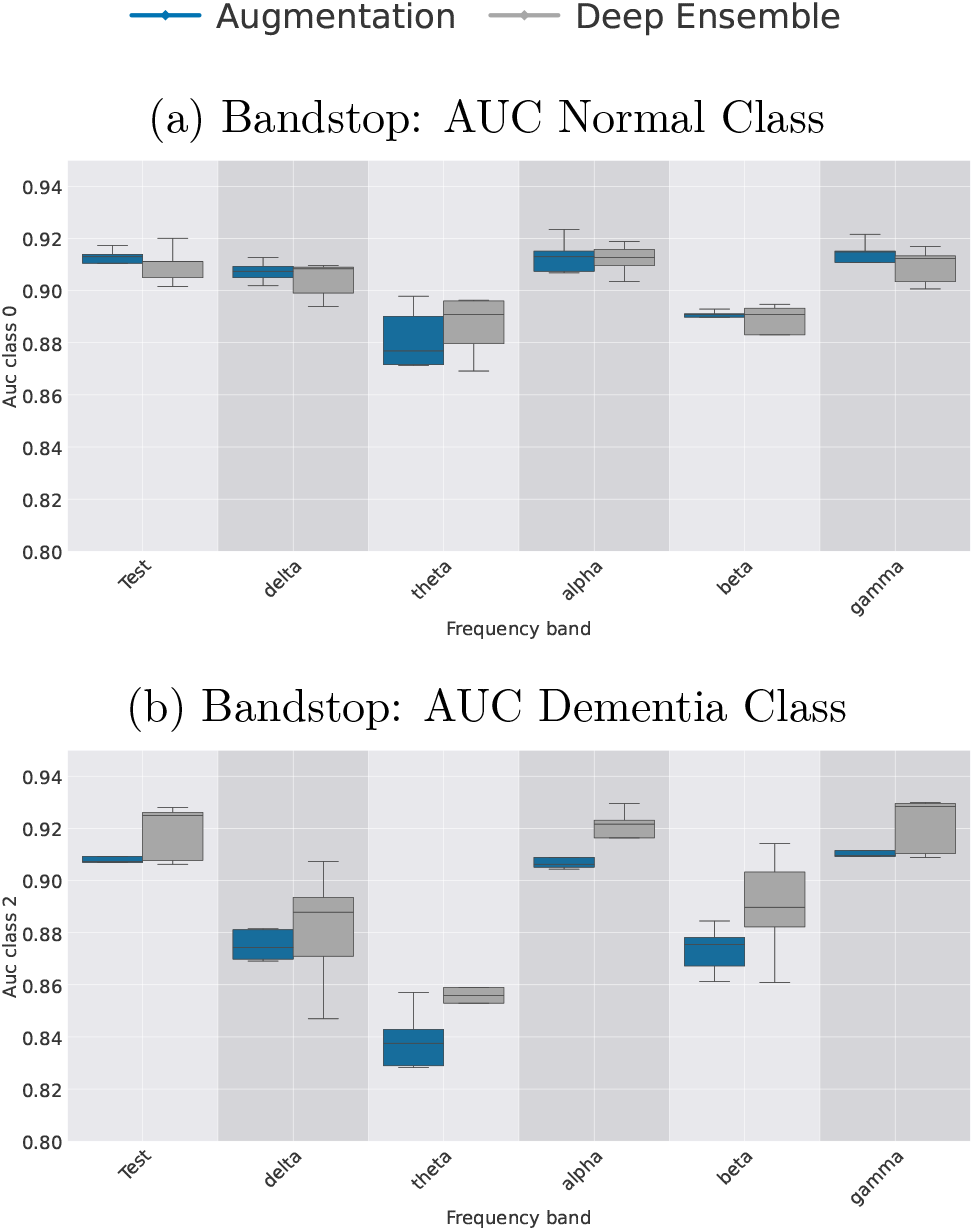
Per-class performance under bandstop shifts. (a) AUC for the Normal class and (b) AUC for the Dementia class, shown as a function of the frequency band removed. Results are shown for the two best-performing ensemble methods, indicated at the top of each plot. The MCI class was excluded due to unstable and uninformative results, and is therefore not displayed.

## 4. Discussion

This study examined how different DL ensemble techniques and MCD classify Normal, MCI and Dementia from EEG and, more importantly, how well their uncertainty and calibration behave under distributional shift. Uncertainty and calibration were evaluated on the in-distribution test set, three OOD datasets and through a series of synthetic dataset shifts designed to represent feature-specific EEG noise. Across these conditions, multi-run ensembles consistently outperformed the other methods in terms of predictive accuracy while also maintaining a meaningful relationship between performance and uncertainty. These findings suggest that multi-run ensembles are better suited for robust and uncertainty-aware EEG classification in the face of data variability.

### 4.1. In-Distribution test

Our best performing multi-run ensembles (Augmentation and Depth) achieved an AUC of 0.84 *±* 0.02 and an accuracy of 68.21% *±* 2.75, falling short of the 0.90 AUC and 74.66% accuracy reported in the original CAUEEG paper [28]. However, the reference model relied on hybrid 1D+2D CNNs, test-time augmentation (which have shown to improve performance [53]), and over 250 million parameters. In contrast, our ensembles used lightweight architectures with only 2.5 million parameters. Other published studies report even higher EEG classification accuracies in similar classification tasks (see [54] for a review), but often on smaller datasets (10-20 times of the current). In addition, many studies use epoch-based classification, evaluating short EEG segments instead of subject-level outcomes. While this artificially increases the sample size, it can produce misleading results if predictions are not aggregated per subject, which is important and considered the correct approach in medical datasets [55]. This epoch-base approach can also introduce data leakage between training and test sets [56, 57].

### 4.2. Comparing performance and uncertainty across test set and OOD

Multi-run ensembles showed consistently better performance over checkpointand samplingbased ensembles in both in-distribution and OOD conditions. They delivered higher predictive performance, particularly for Normal and Dementia cases (AUC ≥ 90), and was associated with lower uncertainty metrics. MCI remained the most difficult class, with an AUC below 0.71, possibly reflecting its clinical heterogeneity and unstable nature [58, 59]. Checkpoint ensembles performed substantially worse across all metrics, suggesting that the limited diversity obtained from a single training trajectory is insufficient for generalisable EEG-based classification. In contrast, MCD, while based on a single model with stochastic dropout, showed intermediate performance, but with wider confidence intervals and greater variability across runs. These differences underscore the importance of ensemble diversity and architectural independence when developing models for complex clinical data.

Multi-run methods also showed higher quality of probabilistic predictions, through Brier scores, on the test set. However, the Brier scores were relatively high (≥ 0.45), and even higher for the checkpoint and sampling methods. These values reflect limited confidence in the model outputs, most likely due to the challenging nature of MCI classification, which generally had a very poor performance, and the ecological and difficult variability in the dataset. In contrast, the ECE values were modest (0.08–0.010), suggesting moderate calibration. However, in several cases, especially for Snapshot and FGE, low ECE coincided with high Brier scores and poor accuracy, indicating underconfident but poorly performing models. This highlights the risk of over-interpreting ECE in isolation and supports the use of Brier score in addition as a more informative measure of uncertainty quality.

Under OOD conditions, differences between ensemble types became more pronounced. Multi-run ensembles kept strong performance in the Militadous cohort, with high recall for Normal cases and near-perfect precision for Dementia. MCD followed a similar trend but with greater variability, while checkpoint ensembles and SWAG degraded substantially, showing wide confidence intervals and reduced precision and recall. In the simpler single-class datasets (MPI, TDBRAIN), multi-run ensembles identified normal cases with near-perfect recall and narrow confidence intervals, confirming their robustness across settings. Age differences between the training dataset (CAUEEG; mean age 71.13) and OOD datasets (Miltiadous: 66.17, MPI: 38.26, TDBRAIN: 29.02) exist. Age was included as an input feature due to its clinical relevance and predictive value, and was regularized during training to reduce overreliance. While prior analyses on the test set suggested that the model’s sensitivity to age was limited, this is not further examined in the current paper. The strong performance on younger OOD cohorts may therefore still partly reflect demographic effects, but assessing this more directly is left for future work.

The uncertainty metrics reflected these shifts. Brier scores increased in the Militadous cohort across all methods, most notably for checkpoint- and sampling-based ensembles, indicating reduced predictive confidence. SWAG was a partial exception, showing lower average Brier scores but much wider confidence intervals, reflecting instability rather than improved reliability. Multi-run ensembles showed smaller increases in Brier score compared to other methods. In the single-class OOD datasets, Brier scores improved, most likely due to task simplicity with only the normal class which the model were generally better to classify, while ECE increased across all models. This divergence illustrates that calibration can degrade even when accuracy improves, reinforcing the importance of interpreting Brier and ECE together. Notably, the relatively low Brier scores in these simpler OOD tasks suggest that the elevated scores observed on the test set are less a sign of model weakness and more a reflection of the in-distribution data’s complexity, particularly in the poorly separated MCI class.

### 4.3. Clinical diagnosis

In addition to the distribution shifts explored in this study, an important source of variability lies in how diagnostic labels are assigned, both within and across datasets. This is particularly relevant when evaluating clinical conditions such as MCI. Diagnostic labels in these domains are functional, and reflect a clinical evaluation of the patient and not definitive biological markers. As such, the diagnostic labelling is subject to observer variability and local protocol differences [60, 61, 62]. These epistemic uncertainties likely contributed to the reduced model performance and elevated uncertainty observed, especially for MCI. This underscores the need for caution when interpreting model results on clinically defined labels.

### 4.4. Dataset shifts

Under distribution shift, we ideally expect model accuracy to decline while predictive uncertainty rises in tandem, allowing downstream systems or clinicians to recognise when predictions become unreliable. Well-calibrated models should maintain meaningful calibration even under shift, not by preserving low error, but by signalling increasing uncertainty as the input distribution diverges [13]. This behaviour enables uncertainty-aware decision making and makes predictive reliability transparent beyond the training domain.

Across the ten dataset shifts, multi-run ensembles consistently demonstrate greater robustness than checkpoint- and sampling-based methods, both in terms of predictive performance and uncertainty estimation. In most shifts, such as amplitude changes, Gaussian noise, and phase shift, performance declines gradually with increasing shift intensity, and uncertainty measures (Brier score and ECE) rise in parallel, especially for the better-performing multi-run models. This alignment suggests that their uncertainty estimates remain informative as conditions degrade. Among the sampling-based methods, MCD most closely resembles the behaviour of multi-run ensembles, showing responsive uncertainty and calibration, but with wider confidence intervals and more pronounced performance degradation. In contrast, SWAG performs inconsistently across shifts, with unstable predictions and wide variability in both performance and uncertainty, limiting its reliability. Under shifts such as timewarp, circular shift, and heavy channel rotation, all models experience sharper performance drops. Even then, Deep and Augmentation ensembles maintain comparatively higher performance. Notably, checkpoint ensembles often show flatter Brier and ECE curves even as performance deteriorates. While flat calibration metrics may appear favourable in isolation, in this context they indicate that the model fails to adjust its confidence in response to increasing error, an undesirable property under distribution shift. In contrast, multi-run ensembles generally scale their uncertainty more appropriately, with both Brier and ECE increasing alongside error. The bandstop and phase shifts revealed frequency specific vulnerabilities, especially in the theta and delta ranges, but again the relative ensemble ranking remains consistent, with multi-run models outperforming others. A closer look at the amplitude and Gaussian noise shifts reveals a distinct pattern: checkpoint ensembles maintain stable AUC, Brier, and ECE values across the full intensity range. This apparent invariance is notable, particularly under extreme conditions where multi-run ensembles begin to degrade. However, this stability appears to be shift-specific, as checkpoint models perform less consistently under other transformations. Overall, these results highlight that multi-run ensembles are not only more accurate but also more reliable under dataset shift, an essential property for deployment in dynamic clinical environments. This aligns with previous research, which has documented the strong performance of such ensembles in different contexts [13]. While their performance declined with increasing shift severity, their uncertainty estimates scaled appropriately, indicating adaptive and informative behaviour. In contrast, checkpoint ensembles often exhibited flat uncertainty curves despite worsening predictions, suggesting a disconnect between confidence and accuracy. This lack of responsiveness may appear robust, but ultimately limits their reliability. This clearly shows that ensemble strategies vary in their response to different types of dataset shift, highlighting the importance of assessing performance and uncertainty together.

### 4.5. Robustness to temporal, amplitude and structural shifts

The dataset shifts explored in this study were constructed by systematically altering specific characteristics of the EEG signal, including temporal distortions, frequency-specific manipulations, amplitude scaling, and structural disruptions. While all models exhibited some vulnerability, the impact of shifts varied substantially. Examining all shifts across all models may reveal which perturbations are most disruptive and which signal properties models consistently depend on, insights that could inform both model development and EEG preprocessing strategies. Amplitude shifts led to steeper performance drops under strong amplification (×8) than under extreme reduction (×0.001), showing that over-amplified signals distort features more severely. Gaussian noise had limited impact at low levels but became damaging as the noise increased, indicating some tolerance to moderate perturbations. Timewarp and circular shift caused abrupt performance decline at the first shift level. For timewarp, this was followed by a gradual decline; for circular shift, performance remained relatively stable after the initial drop. This pattern may reflect edge effects introduced when appending the time signal to itself, and possibly the translational invariance of CNNs, which may reduce the sensitivity to the degree of temporal offset once the signal is disrupted. Slow drifts also degraded performance, highlighting the importance of removing low-frequency components, e.g., through high-pass filtering. Structural distortions had varied effects. In interpolation, performance remained stable until more than 75% of the channels were removed, which may have been due to central channels Cz and Pz being consistently preserved. Beyond that threshold, the accuracy dropped sharply. Channel rotation led to a steady decline, underscoring the importance of preserving spatial structure and consistent channel ordering between training and testing.

### 4.6. Frequency band importance and clinical alignment

Bandstop shifts revealed that the delta, theta, and beta bands are particularly informative for this prediction task, with their removal causing measurable performance drops. A perclass analysis provided more detail: for the Normal class, removing either theta or beta led to a notable drop in AUC, suggesting both bands contribute to the model’s ability to distinguish normal EEG. In contrast, dementia classification showed the strongest sensitivity to theta-band removal, with a more modest impact from delta and beta.

Although this study primarily focused on method development, some findings align with established EEG markers in dementia. Specifically, the model’s reliance on theta, delta, and beta bands reflects well-documented spectral changes in Alzheimer’s disease, where increased slow-wave activity and reduced beta power are linked to cortical disconnection and cognitive decline [63, 64, 65].

In contrast, the alpha band’s contribution was minimal, a surprising finding given its well-documented importance in clinical literature. This conclusion is supported by two distinct lines of evidence: the direct removal of the alpha band via bandstop and the separate alpha peak-shift experiment both resulted in only a gradual performance decline. Although the peak-shift manipulation was less controlled, using random, bidirectional, and channel-specific changes, the consistency of its outcome with the more direct bandstop analysis lends weight to the conclusion. This limited role of alpha is likely attributable to the data originating from a mixed pre-task period (including both eyes-open and eyesclosed states) and not from a dedicated eyesclosed resting state, where alpha rhythms are most prominent. Thus, in this limited sample, the alpha band did not emerge as a significant predictive feature for the implemented models.

### 4.7. Limitations and future work

Our study has several limitations that provide avenues for future research. First, while the primary findings are based on a test set of 117 subjects, the robustness of our approach was further stressed against several OOD datasets. Although this represents an extensive evaluation, the conclusions still require validation on larger and more diverse multi-centre samples to establish broader generalisability. Second, our OOD evaluation focused on the Normal and Dementia classes and did not include cases of MCI. Assessing model robustness for this group is a critical future direction; however, it is crucial to recognize that this is complicated by the known clinical heterogeneity of MCI. Future work involving MCI must carefully account for variability in diagnostic criteria between datasets, which acts as a major confounding factor in the absence of consistent biological markers. These clinical inconsistencies, combined with technical differences in EEG recording hardware and preprocessing pipelines across sites, pose a significant challenge to model generalisability. Furthermore, we employed a deliberately minimal preprocessing pipeline to assess robustness in a manner that better reflects scalable deployment. While this choice may have constrained optimal model performance, it offered a test of real-world generalisability. Finally, the synthetic shifts used here for evaluation could themselves be leveraged as a data augmentation strategy during training, a promising approach to proactively enhance model robustness and clinical translation.

## 5. Conclusion

Multi-run ensembles, such as deep ensemble, consistently delivered the best performance and most reliable uncertainty estimates across test, OOD, and shifted EEG data. Their ability to align uncertainty closely with changes in predictive performance under distribution shifts makes them the most promising approach for robust, uncertainty-aware EEG classification. Together, these findings underscore the value of ensemble-based uncertainty modelling as a foundation for building trustworthy and clinically deployable EEG classification systems.

## Project Information and Contributions

### Funding

This project has received funding from the European Union’s Horizon 2020 research and innovation programme under grant agreement No 964220. This publication reflects views of the authors and the European Commission is not responsible for any use that may be made of the information it contains.

### Author Contribution

**Mats Tveter:** Concept, Software, Investigation, Methodology, Validation, Formal Analysis, Visualisation, Writing - Original Draft, Writing - Review & Editing **Thomas Tveitstøl:** Visualisation, Writing - Review & Editing **Christoffer Hatlestad-Hall:** Supervision, Writing - Review & Editing **Hugo L. Hammer:** Supervision, Visualisation, Writing - Review & Editing **Ira R. J. Hebold Haraldsen:** Project administration, Writing - Review & Editing, Funding acquisition, Resources

### Data Availability

The dataset is accessible as open source or upon request. For further details, consult the studies: CAUEEG [28], Militadous [31] MPI, and TDBRAIN [33].

Code for all experiments and models can be found on GitHub∥.

## Footnotes

‡ For practical reasons, we use the name of the first author to refer to this dataset throughout the paper

